# Effects of single mutations on protein stability are Gaussian distributed

**DOI:** 10.1101/2019.12.17.879536

**Authors:** R. M. Razban, E. I. Shakhnovich

## Abstract

**Abstract:** The distribution of protein stability effects is known to be well-approximated by a Gaussian distribution from previous empirical fits. Starting from first-principles statistical mechanics, we more rigorously motivate this empirical observation by deriving per residue protein stability effects to be Gaussian. Our derivation requires the number of amino acids to be large, which is satisfied by the standard set of 20 amino acids found in nature. No assumption is needed on the protein length or the number of residues in close proximity in space, in contrast to previous applications of the central limit theorem to protein energetics. We support our derivation results with computational and experimental data on mutant protein stabilities across all types of protein residues.

**Statement of Significance:** Defining the distribution of single mutant stability effects (ΔΔGs) is the first step in modeling the role protein stability plays in evolution. Although empirical fits have been made to elucidate its form, a complete theoretical understanding of ΔΔG distributions is lacking. Here, we derive how a simple Gaussian form can arise, while still including the intricacies of protein sequence and structure. We backup our derivation with previously released computational and experimental ΔΔGs.

## Introduction

The overall stability of a protein is influenced by its residues and their interactions (1). Any change to the wildtype sequence, even a single mutation, has the potential to drastically affect stability because proteins are marginally stable (2, 3). Since mutations fix readily across evolution (4, 5), careful consideration must be made whether a certain mutation fixing in the population is the result of that mutation’s stability effect. Unfolded proteins may no longer carry out their function in the cell (6) or may form toxic aggregates (7, 8). One of the first steps in modeling stability constraints requires defining the distribution of single mutant stability effects (ΔΔGs)^1^ (3, 9–13).

Zeldovich et al. (2007) took experimental ΔΔG data from different proteins deposited in the Protein Thermodynamic database (14) and found the distribution to be well-approximated by a Gaussian. Although limited by experimental data, Zeldovich et al. (2007)’s results seems to extend to individual proteins. Tokuriki et al. (2007) found a universal Gaussian form for all possible ΔΔGs calculated for a given protein using FoldX (16), a fast and relatively accurate computational software that requires a Protein Data Bank (PDB) (17) structure as an input. Faure and Koonin (2015) extended upon Tokuriki et al. (2007)’s study of 16 natural proteins by using FoldX to calculate ΔΔGs for all proteins with PDB structures in 5 organisms’ proteomes. They also found a universal Gaussian form for proteins of mesophilic proteomes (18).

In this paper, we apply first principles statistical mechanics to show that per residue ΔΔG distributions can by accurately approximated by Gaussians with means and variances shaped by their local (in space) sequence and structure. Our derivation elucidates that the large number of amino acid types is responsible for the Gaussian being a good approximation, unlike previous arguments in the field which relied on the large number of pairwise contact energies making up protein energy (10, 19). Indeed, we show that our theory applies almost equally as well to those residues with fewer contacts for Tokuriki et al. (2007)’s computational data and for newly released experimental data from a study that quantified almost all ΔΔGs of a protein G domain (20).

## Derivation

We solve for the distribution of energy effects for a given residue by first applying a continuous spin approximation to obtain the partition function of energy effects (21) and then employing an inverse Laplace transform over inverse temperature to proceed from the partition function to the density of states (22), also known as the distribution of energy effects.

### Protein energy and formalism

The energy of the protein (E) is defined as the sum over individual contact energies,

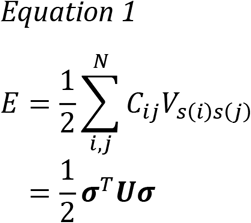

N is the number of amino acids in the protein. **C** is a N*N contact matrix whose elements can only take values of 1 (two residues spatially close in proximity, or in contact) or 0 (not in contact). **V** is a M*M interaction matrix, where M is the number of amino acid types (M = 20 standard amino acids), and an element of **V** is the pairwise energy of two specific amino acids when they are in contact. The variable s refers to the wildtype sequence; s(i) refers to the amino acid of residue (i). **σ** is a NM dimensional vector, where M is the number of amino acid types. **U** = **C** ⊗ **V** and is a NM ∗ NM dimensional matrix. Integrals will be simpler to perform with matrices, thus the rewrite in the second line of Equation 1. Each N block of σ has M elements, in which one element is equal to one while the other M − 1 elements are equal to zero. The index of the non-zero element sets the amino acid type.

Since we are interested in single mutant effects at a residue (i), we want ΔE_(i)_ = E_mut,(i)_ − E_wt_. First, we rewrite E in terms of the following matrix blocks,

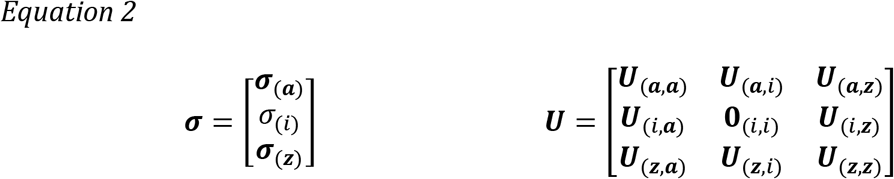

**σ_(a)_** corresponds to all σs numbered before residue (i); **σ_(z)_**, after residue (i). **U**_(i,i)_ = **0**_(i,i)_ is an M ∗ M dimensional zero matrix because the contact matrix has zeros along the diagonal. In other words, residues cannot be in contact with themselves. In addition, **U** is symmetric because **C** and **V** are symmetric by definition.

We commence to write the energy (Equation 1) in terms of the defined matrix blocks,

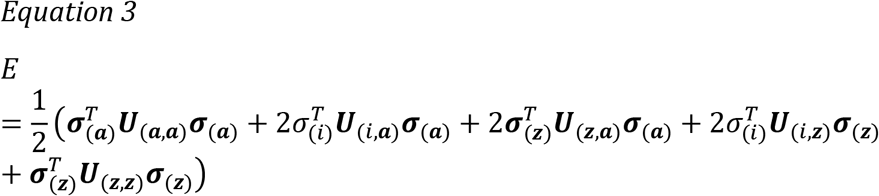

Using Equation 3, ΔE_(i)_ can be compactly expressed as,

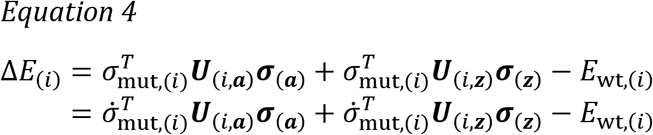

Terms not involving (i) cancel because they are the same across mutant and wildtype. We also reduce σ_(i)_ to only include amino acids that are non-wildtype in the second line of Equation 4; 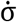 refers to an M-1 dimensional vector that does not include the wildtype amino acid. The same notation applies for **U**.

For future convenience in evaluating the partition function, let’s define 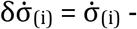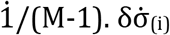 has the nice property of having a length of 0 when referring to an amino acid.

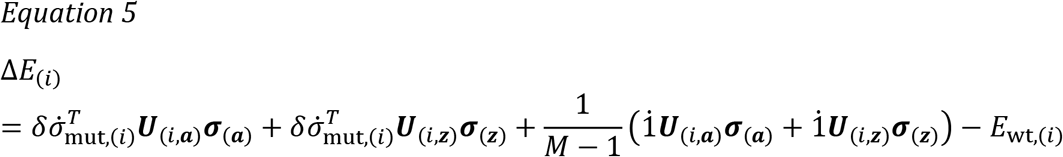

### Partition function

The partition function for one residue mutating can be approximately calculated by integrating each mutant term in 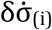 over all space and picking out terms that correspond to a certain amino acid type with a Dirac delta function δ(x).

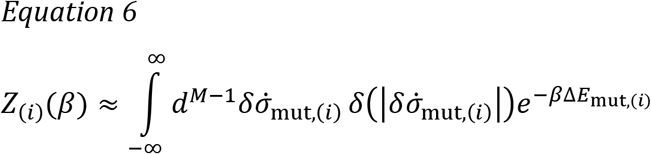

This is the form of the continuous spin approximation that England and Shakhnovich (2003) also employed, except here we integrate over all mutant amino acids for residue (i). Equation 6 is an approximation because the Dirac delta function as written does not generally correspond to the aforementioned condition that only one element of 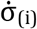 is equal to 1, while all other M − 2 elements are zero. Multiple terms can have values less than 1, such that the length of 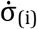 (denoted as 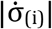) equals to 1, or equivalently 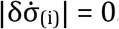. All amino acids and residues are equally affected; thus, we assume qualitative results are retained despite including those extra terms.

To make more progress, the delta function can be well-approximated by a Gaussian distribution when the variance approaches zero (23). This approximation is key to making the integral soluble and limiting terms to quadratic in 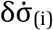. The variance can be expressed as,

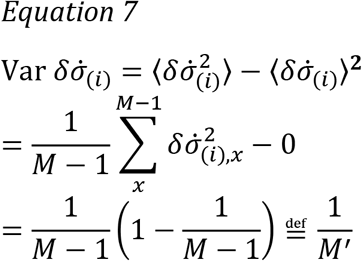

Thus, our Gaussian approximated delta function leads Equation 6 to look like,

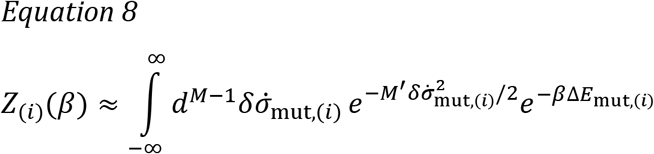

We can now begin solving the integral. First, we insert the expression for ΔE_(i)_ (Equation 5) into Equation 8 and pull out constants in front of the integral. Then we complete the square of the exponent in the second line.

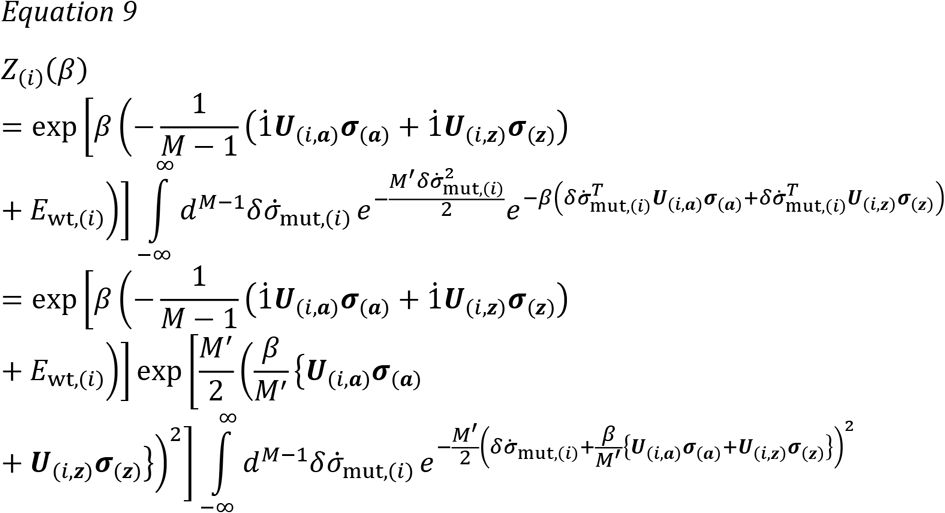

The multidimensional integral represents M −1 identical Gaussian integrals and can be easily solved to be a constant independent of β.

Before moving on and obtaining the density of states, let’s do two things. We define a reduced partition function z(β) = Z(β)/Z(0), since constants will eventually cancel out when calculating the density of states and ensuring that it is normalized. Let’s also revert matrices back to the more intuitive summation notation.

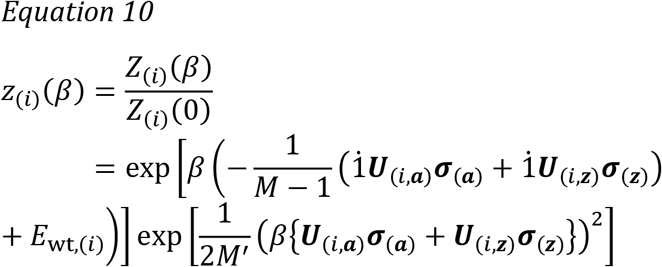

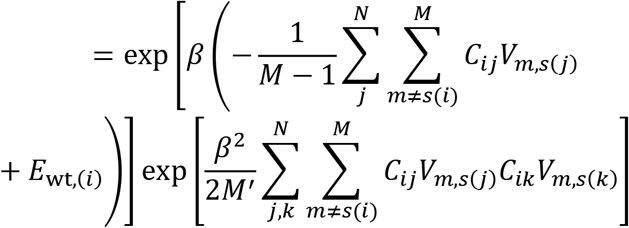

### Density of states

The density of states (n) is related to the partition function by an inverse Laplace transform (22).

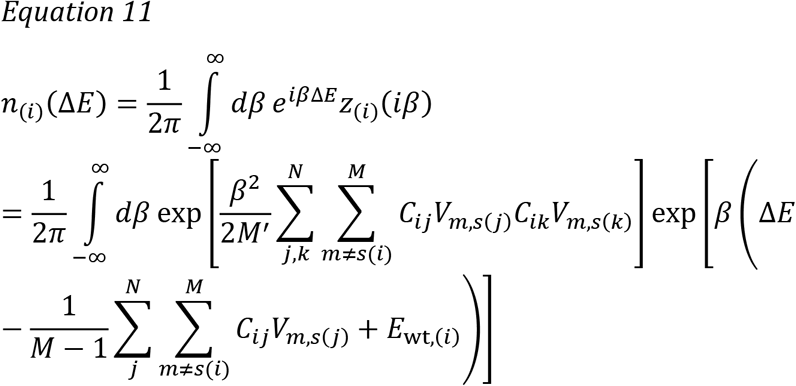

The integral can be solved by completing the square. Skipping steps, it can be shown that the following Gaussian distribution (G(mean, variance)) is exactly obtained.

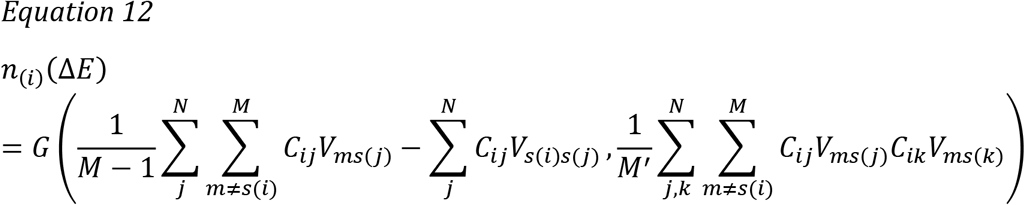

The expressions for the mean and variance of the Gaussian in Equation 12 are intuitive. The mean is <ΔE_(i)_> and the variance is <ΔE_(i)_^2^> if M’ = M−1. This is what we would expect if we were to fit a Gaussian to data whose energy is determined by Equation 1, except that the variance would be <ΔE_(i)_^2^> − <ΔE_(i)_>^2^. Some inaccuracy is reasonable because we employed a continuous spin approximation (Equation 6) and approximated the delta function as a Gaussian (Equation 8).

Our derivation found a distribution for ΔE, not ΔΔG. We assume that the unfolded state and entropic effects are less important and can be accounted by properly fitting the interaction matrix, thus not changing the final form of the distribution. In other words, we assume n_(i)_(ΔE)~n_(i)_(ΔΔG).

## Bioinformatics

### Computational ΔΔGs

To test whether per residue ΔΔG distributions are Gaussian (Equation 12), we first organize ΔΔGs from Tokuriki et al. (2007) into groups of 19 ΔΔGs belonging to individual residues to obtain per residue ΔΔG distributions (Materials and Methods). We then run the Shapiro-Wilk test (24) to analyze how well a Gaussian distribution fits the 19 ΔΔGs, with a mean and variance calculated from the data. Our null hypothesis is that the distribution is a Gaussian. As the Shapiro-Wilk test statistic decreases and the p-value subsequently decreases, we can be more confident that the distribution is not Gaussian, and our null hypothesis is wrong (Materials and Methods). For example, in Figure S1 we run the Shapiro-Wilk test on the first residue of dihydrofolate reductase (DHFR) and find that the test statistic is small enough that we can reject the null hypothesis with a p-value threshold of 0.05. In other words, the 19 ΔΔGs pertaining to the first residue of DHFR is not Gaussian distributed, at odds with our prediction (further discussed below).

Running the Shapiro-Wilk test for all residues from the 16 natural proteins in Tokuriki et al. (2007), we find that a majority of residues have ΔΔG distributions indistinguishable from a Gaussian (Figure 1). We further investigate the 10-25% of residues that do reject the null hypothesis at a p-value threshold of 0.05, like the first residue of DHFR discussed above. In Figure 2, we show box plots of contacts for residues that reject the null and for all residues. For 11 proteins, the median number of contacts for residues that reject the null is less than the median number of contacts for all residues. For five proteins, the medians are equivalent; and for one protein, Ribonuclease A (RNaA), the median number of contacts for residues that reject the null is greater than the median number of contacts for all residues. However, a more detailed analysis comparing the distributions of the number of contacts using the Kolmogorov-Smirnov test (25), yields only human carbonic anhydrase II (CAII) to have significantly different contact distributions between those residues that reject the null and all residues (p-value = 0.037). The lack of a structural signature of those residues that reject the null makes sense in light of our theory, which does not require an assumptions on the number of contacts.

**Figure 1:**
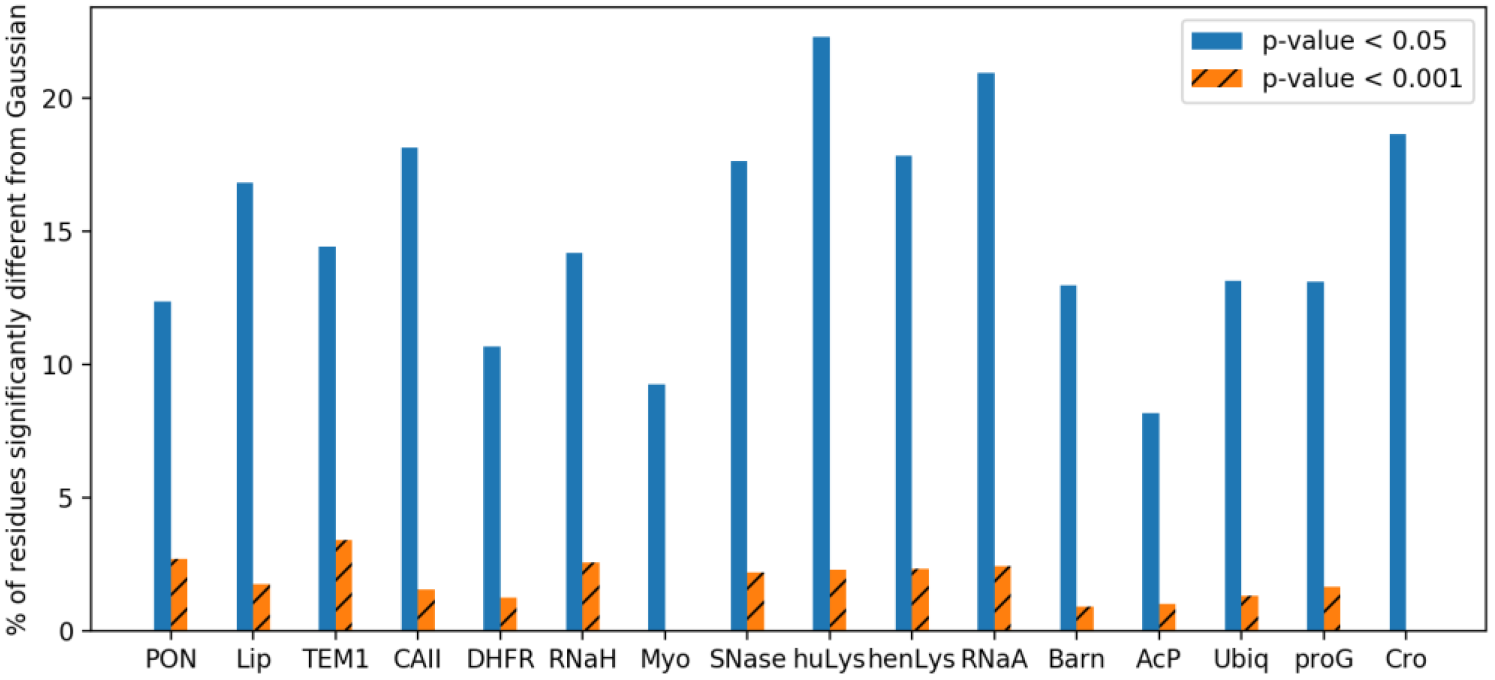
Relatively few residues per protein fail to match our theoretical prediction that per residue ΔΔGs are Gaussian distributed according to the Shapiro-Wilk test. Solid bar plots correspond to residues that reject the null at a p-value threshold of 0.05; hatched, reject the null at a p-value of 0.001. For protein acronyms please see Table 1 in Materials and Methods.

**Figure 2:**
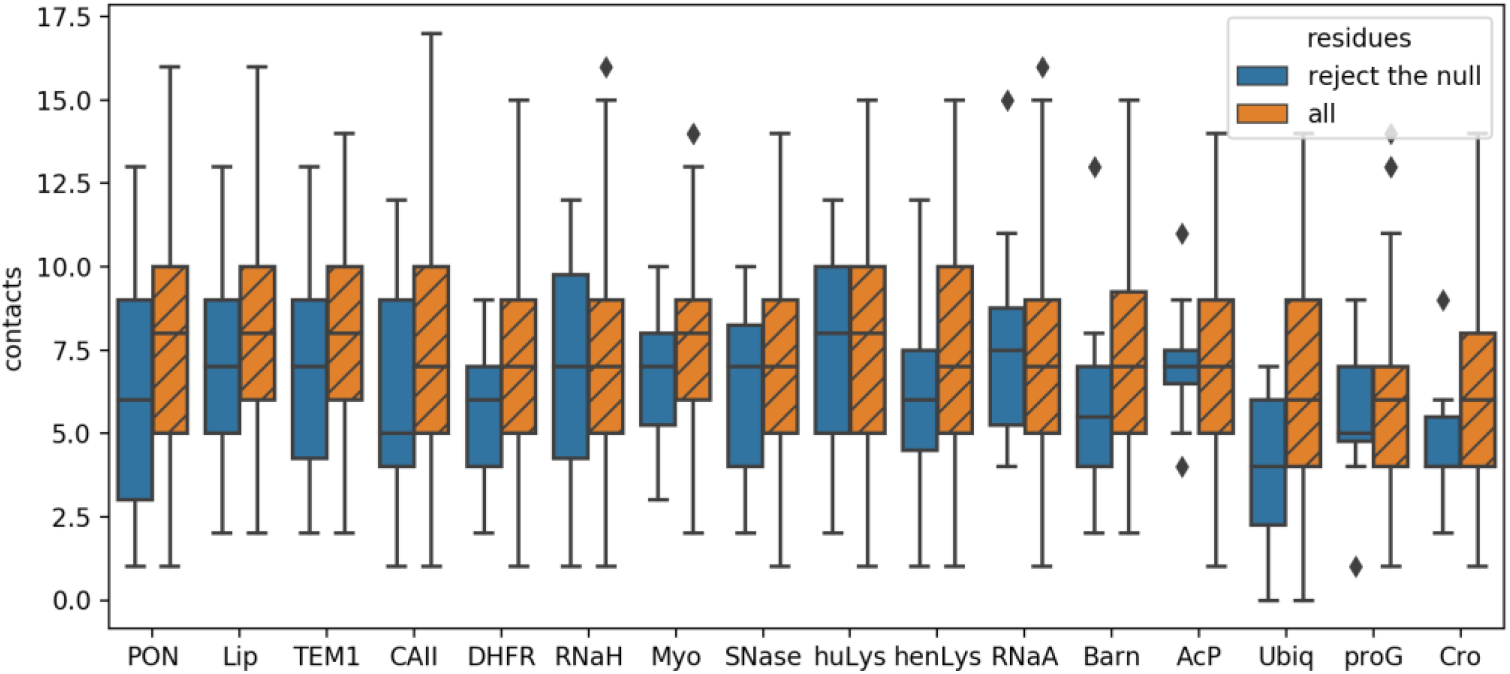
The distribution of contacts for residues found to reject the null hypothesis in Figure 1 at a p-value threshold of 0.05 (solid box plots) is similar to that of all protein residues (hatched box plots). Horizontal black lines correspond to medians; diamonds correspond to outliers.

**Table 1:** Proteins whose ΔΔGs are studied. All protein ΔΔGs, except Gβ1, originate from Tokuriki et al. (2007). Gβ1 ΔΔGs were experimentally measured by Nisthal et al. (2019). Not all of Gβ1 ΔΔGs were measured because of experimental constraints.

| Protein name | acronym | PDB | length | organism |
| --- | --- | --- | --- | --- |
| Recombinant serum paraoxonase | PON | 1v04 | 332 | *Oryctolagus cuniculus* |
| Lipase | Lip | 1ex9 | 285 | *Pseudomonas aeruginosa* |
| β-Lactamase | TEM1 | 1btl | 263 | *Escherichia coli* |
| Human carbonic anhydrase II | CAII | 1lug | 259 | *Homo sapiens* |
| Dihydrofolate reductase | DHFR | 1rx2 | 159 | *Escherichia coli* |
| Ribonuclease H | RNaH | 2rn2 | 155 | *Escherichia coli* |
| Myoglobin | Myo | 1a6k | 151 | *Physeter macrocephalus* |
| Staphylococcus nuclease | SNase | 1stn | 136 | *Staphylococcus aureus* |
| Human lysozyme | huLys | 1rex | 130 | *Homo sapiens* |
| Hen lysozyme | henLys | 1dpx | 129 | *Gallus gallus* |
| Ribonuclease A | RNaA | 1fs3 | 124 | *Bos taurus* |
| Barnase | Barn | 1a2p | 108 | *Bacillus amyloliquefaciens* |
| Acylphosphatase | AcP | 2acy | 98 | *Bos taurus* |
| Ubiquitin | Ubiqu | 1ubq | 76 | *Homo sapiens* |
| Protein G, IgG-binding domain | proG | 2igd | 61 | *Streptococcus sp. group G* |
| Cro repressor | Cro | 1orc | 59 | *Escherichia phage lambda* |
| Protein G, β1 domain | Gβ1 | 1pga | 56 | *Streptococcus sp. group G* |

With a more stringent p-value threshold of 0.001, no protein rejects the null hypothesis for more than 4% of its residues (hatched bar plots in Figure 1). We can argue that a more stringent p-value threshold is necessary to address the multiple testing problem (26).

### Experimental ΔΔGs

A major caveat of the previous subsection is that ΔΔGs are generated from FoldX. As noted in the Introduction, exhaustive studies of protein ΔΔG distributions have been limited to computational calculations because the number of experimental measurements is too large to complete in a reasonable timeframe. Recently, Nisthal (2012) made exhaustive ΔΔG measurements feasible by developing an experimental protocol utilizing automated robotics for protein purification and chemical denaturant curve measurements. With this protocol, Nisthal et al. (2019) quantitatively measured the ΔΔGs of almost all single mutants (78%) of the β1 domain of Streptococcal protein G (Gβ1).

We find that per residue ΔΔG measurements are mostly Gaussian distributed, but at a ratio on the high end of those found in Figure 1. Including all Gβ1 residues with at least three measured ΔΔGs (54 out of 56 total residues), 13 (24%) residues reject the null hypothesis at a p-value threshold of 0.05; 6 (11%), at a p-value of 0.001 (*Figure 3*). We tested whether having few ΔΔGs measured for some residues explained the relatively high percentages of null hypotheses rejected, however, this was not the case. When only analyzing residues with the maximal number of 17 ΔΔGs measured (29 out of 56 total residues), 11 residues still reject the null at a p-value threshold of 0.05; 5, at a p-value of 0.001 (*Figure 3*). The contact distribution of those 11 residues that reject the null does not matter either. We find that the contact distribution is indistinguishable with those of all residues according to the Kolmogorov-Smirnov test (p-value = 0.40).

**Figure 3:**
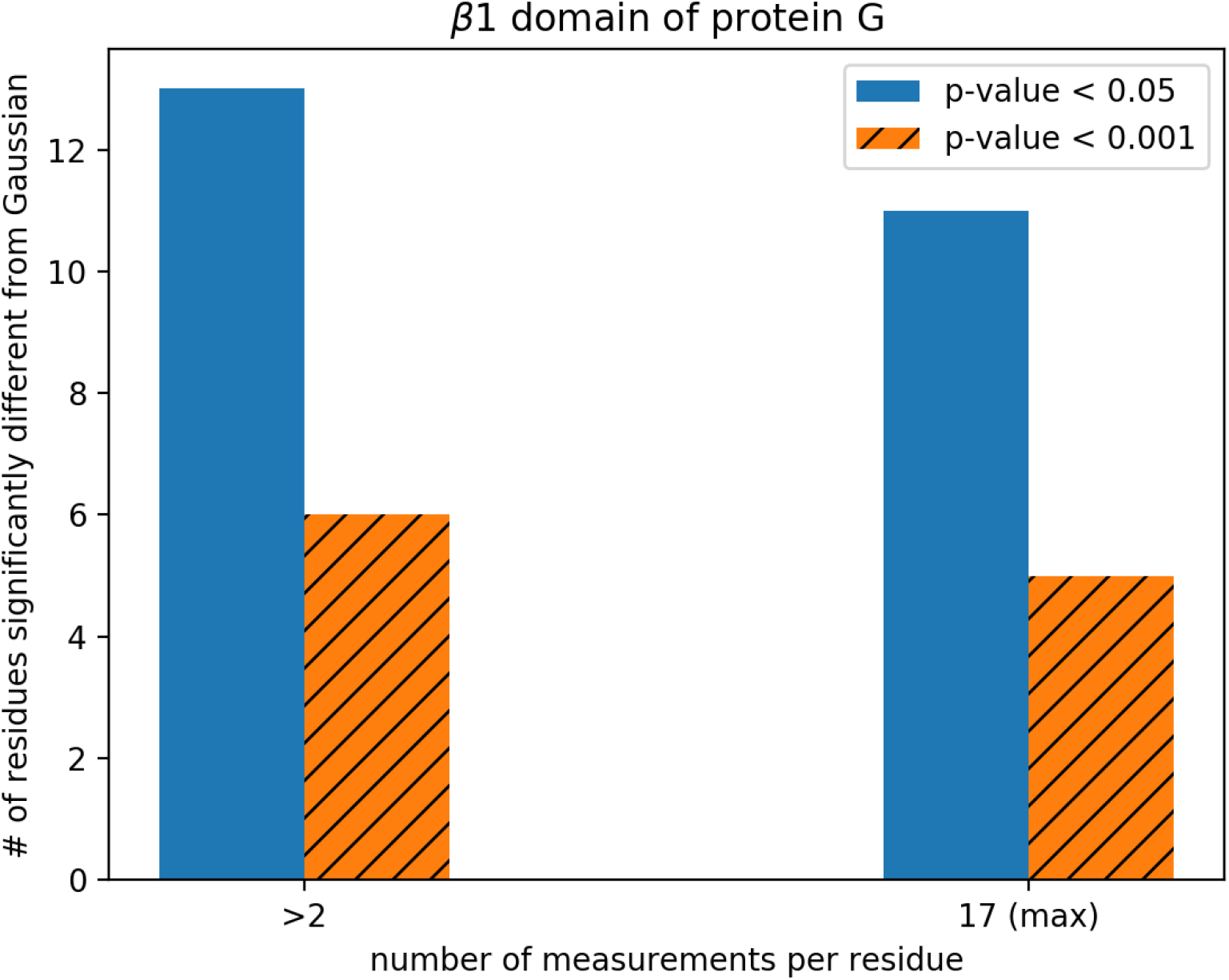
Relatively few residues of β1 domain of Streptococcal protein G (56 total residues) fail to match our theoretical prediction that per residue ΔΔGs are Gaussian distributed according to the Shapiro-Wilk test. Solid bar plots correspond to residues that reject the null at a p-value threshold of 0.05; hatched, reject the null at a p-value of 0.001. Residues that reject the null mostly have the maximal number of mutants measured in the study (right two-most bar plots).

## Discussion

Employing a previously established theoretical framework (21), we derive from first principles statistical mechanics that per residue ΔΔGs are Gaussian distributed. Our theory yields a Gaussian while still considering the intricacies of the protein structure and unique interactions of amino acid types. We verify with computational and experimental data that per residue ΔΔGs are Gaussian distributed for most residues (Figure 1 and Figure 3).

The finding that per residue ΔΔGs are Gaussian distributed may not seem surprising in light of the central limit theorem. However, the central limit theorem applies not because individual contact energies can be treated as random variables, but because each residue has 19 possible mutant amino acid identities (Equation 8). This yields Gaussian distributions for all residues, regardless of its number of native contacts (Figure 2).

The mathematical framework we employed closely follows that of England and Shakhnovich (2003). They derived from first principles an expression for the number of sequences that folds into a given structure, called protein designability. Unlike here where we integrate over all mutant amino acid types for a given residue (Equation 6), England and Shakhnovich (2003) integrated over all amino acid types across all residues to obtain the partition function. Interestingly, when applying an inverse Laplace transform to obtain the density of states, England and Shakhnovich (2003) cannot exactly obtain a Gaussian. Within the theory, a Gaussian distribution is only exactly obtained for per residue mutants. However, when considering all possible mutants at once (not just single mutants), a Gaussian distribution can only be approximated.

Our derivation falls short from generally showing that the overall ΔE distribution of all single mutants (n(ΔE)) is Gaussian. By definition, n(ΔE) is the weighted sum of all N Gaussians from Equation 12 because it corresponds to sequentially picking a single mutant out of N residues, i.e. a statistical OR operation.

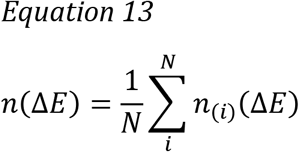

We are not aware of an approach using first principles to reduce the mixture of N Gaussians in Equation 13. ^2^ However, we can bioinformatically demonstrate that our N Gaussian model reduces to a bi-Gaussian (Figure S2 and Figure S3). Therefore, a more tractable Gaussian mixture model with fewer Gaussians than the exact N Gaussian model elucidated here can be employed with little loss in accuracy.

Future studies can use Equation 12 to fit a 20*20 interaction matrix to yield accurate means and variances of per residue ΔΔG distributions for experimental data. A 20*20 interaction matrix is too simple to generally capture protein folding (28), however for single mutant ΔΔGs that level of detail may suffice.

## Acknowledgements

We are grateful to Dan Tawfik and Alex Nisthal for fruitful email exchanges. We would also like to thank Bharat Adkar and Mobolaji Williams for useful comments on this manuscript.

This work is supported by the National Institutes of Health (grant number GM067680).

## Author Contributions

RMR and EIS designed research, analyzed data, wrote the manuscript. RMR performed the research.

## Materials and Methods

The computational data for the 16 natural proteins studied in Tokuriki et al. (2007) was kindly provided by Dan Tawfik. The full names of the 16 proteins and the acronyms used in this text, along with their lengths and endogenous organisms can be found in Table 1. The lone protein presented with experimental ΔΔG data, the β1 domain of protein G, is also included at the end of Table 1. The experimental data was kindly provided by Alex Nisthal. We adopt the convention that a more negative ΔG equates to a more stable protein, thus ΔΔG = ΔG^mut^ − ΔG^wt^ < 0 is a stabilizing mutation.

We employ the Shapiro-Wilk (S-W) test to determine whether per residue ΔΔGs are Gaussian distributed. S-W can only be applied for normality testing and has been shown to be the most powerful in this regards (29). If the test-statistic, which ranges from 0 to 1, approaches zero and has an appropriately minuscule p-value, then we can reject the null that our data fits a Gaussian distribution.

We calculated the number of contacts for each residue from in-house scripts analyzing corresponding PDB structures (Table 1). A two-sample Kolmogorov-Smirnov (K-S) test is employed to determine whether our contact distributions between residues that reject the null and all residues differ.

For Figure S2 and Figure S3, the K-S test is employed to determine whether our N-Gaussian model or a bi-Gaussian fit, matches the distribution from the ΔΔG data. We carry out a two-sample K-S test to compare binned data to each other. All binning is performed exactly as done in Tokuriki et al. (2007); ΔΔGs are parsed in 1 kcal/mol bins from −10 kcal/mol to 15 kcal/mol. Calculated FoldX ΔΔGs beyond those limits are set to the corresponding binning limit.

Scripts used to analyze data from Tokuriki et al. (2007) and Nisthal et al. (2019) can be found on GitHub (github.com/rrazban/ddG_distribution).

## Footnotes

1 In this text, stating ‘ΔΔGs’ always refers more broadly to all possible single mutant ΔΔGs.

2 Notice that we are summing normal distributions and not random variables that are normally distributed. The sum of two normally distributed random variables is normally distributed; however, it does not generally hold that the sum of two normal distributions is another normal distribution (30).

